# Design of an Integrated Microvascularised Human Skin-on-a-Chip Tissue Equivalent Model

**DOI:** 10.1101/2022.04.06.487276

**Authors:** Christian F. E. Jones, Stefania di Ciò, John Connelly, Julien Gautrot

## Abstract

Tissue engineered skin constructs have been under development since the 1980s as a replacement for human skin tissues and animal models for therapeutics and cosmetic testing. These have evolved from simple single cell-assays to increasingly complex models with integrated dermal equivalents and multiple cell types including a dermis, epidermis and vasculature. The development of micro-engineered platforms and biomaterials has enabled scientists to better recreate and capture the tissue microenvironment in vitro, including the vascularization of tissue models and their integration into microfluidic chips. However, to date, microvascularised human skin equivalents in a microfluidic context have not been reported. Here we present the design of a novel skin-on-a-chip model integrating human derived primary and immortalized cells in a full thickness skin equivalent. The model is housed in a microfluidic device, in which a microvasculature was previously established. We characterize the impact of our chip design on the quality of the microvascular networks formed and evidence that this enables the formation of more homogenous networks. We developed a methodology to harvest tissues from embedded chips, after 14 days of culture, and characterize the impact of culture conditions and vascularization (including with pericyte co-cultures) on the stratification of the epidermis in the resulting skin equivalents. Our results indicate that vascularization enhances stratification and differentiation (thickness, architecture and expression of terminal differentiation markers such as involucrin and transglutaminase 1), allowing formation of more mature skin equivalents in microfluidic chips. The skin-on-a-chip tissue equivalents developed, thanks to their realistic microvasculature, may find application for the testing efficacy and safety of therapeutics delivered systemically, in a human context.

## 1 Introduction

Our ability to recreate more complex tissue structure and function has considerable improved. As the fields of micro-engineering and biomaterials science developed, it has become increasingly clear that recreating some of the 3D architecture of tissues may be essential to reproduce the cell microenvironment and direct cell phenotype and tissue structure or function. Dating back from the 80s, micro-engineered platforms allowed the control of single cell adhesion and patterning, in turn allowing to control their phenotype (Watt et al., 1988, Chen et al., 1997, Connelly et al., 2010, Tan et al., 2013). More recently, the formation of clusters allowing to capture more complex behaviors, including cell segregation, motility, sprouting and differentiation have been developed (Nelson et al., 2006, Ruiz and Chen, 2008, Gautrot et al., *2012,* Costa et al., 2014). However, these platforms remained inherently based on 2D models, certainly relevant to cells forming 2D assemblies such as epithelia, but clearly limiting the creation of more complex architectures. Although interesting strategies were proposed to bridge the 2D and 3D world (Bosch-Fortea et al., 2019), making use of 2D patterns to guide cell-assembly in a 3D matrix, our ability to micro-engineer 3D tissues required the development of novel platforms.

The field of skin biology was certainly the first to see striking progress in the development of complex hierarchical tissue structures, with stratified differentiated skin equivalents of human tissues forming from epidermal keratinocytes assembling as monolayers at the surface of hydrogels encapsulating fibroblasts (Shahabeddin et al., 1990, Stark et al., 2006). This allowed not only the recreation of well-differentiated human skin equivalents, but also allowed capturing some of the skin barrier functions, from trans-epithelial resistance to the diffusion of therapeutics and nanomaterials (Ackermann et al., 2010, Asbill et al., 2000, Bellas et al., 2012, Mazlyzam et al., 2007). These include models which have integrated other *in vivo* skin structures and cell types including the hair follicle (Langan et al., 2015) and melanocytes (Li et al., 2011, Liu et al., 2007). More recently, a broad range of tissues have beneficiated from advances in stem cell biology, spheroid and organoid models and progress in biomaterials and microfabrication. This ranges from the formation of functional muscle and lung tissue models (Huh et al., 2010, Ronaldson-Bouchard et al., 2018, Sakar et al., 2016) to the development of complex organoid models, for example of gut, skin and brain tissues (Lancaster et al., 2013, Lee et al., 2020).

As tissues grew in complexity and size, vascularization became an important hurdle, either to their further development or growth, or simply their long term survival and applicability (Baish et al., 2011, Coloma et al., 1997, Colton, 1995). In addition, recapitulating the complex architecture and function of microvascularised tissues will be important to the accurate prediction of therapeutics safety and efficacy (Loskill et al., 2021) for example mimicking systemic delivery to targeted tissues. To address this issue, a range of tissue models have been placed in parallel with endothelialised structures, therefore allowing to recreate the endothelium-tissue interface. This has in particular be exploited in a broad range of tissue-on-chip models, using porous membranes to separate an endothelial compartment from a tissue compartment (Hassell et al., 2017, Huh et al., 2010, Kilic et al., 2016, Kim and Ingber, 2013). In addition to simply structuring tissues, this approach enables the mechanical stimulation of resulting tissues, an important factor regulating cell and tissue biology and response to pathogen or during disease progression. In order to recreate more functional models of vessels, a range of platforms have been engineered to allow the endothelialization of micro-capillaries and microchannels, allowing to study phenomena such as sprouting and angiogenesis in more realistic scenarios (Liu et al., 2021, Menon et al., 2017, Trappmann et al., 2017).

Increasingly, microvascularised models have become popular, as enabling to accurately capture the architecture and function of microvascular networks perfusing tissues (Polacheck et al., 2014). Early work from Kamm and co-workers and Jeon et al. showed that microvascular networks can conveniently be assembled and maintained in microfluidic chips, allowing the formation of perfusable capillaries (Galie et al., 2014, Kim et al., 2013). Such models can be derived through angiogenesis, from endothelialised microchannels interfaces with hydrogel compartments separated by micro-posts, or by direct assembly of networks via vasculogenesis, therefore enabling faster formation of functional structures. Subsequently, hydrogel compartment were loaded with stem cells and disease cell lines in order to recreate vascularized tissue models (Chen et al., 2017, Jeon et al., 2015, Kim et al., 2015a). Another attractive alternative proposed consists in placing spheroids, and in some cases organoids within wells designed within the hydrogel compartment, therefore enabling the vascularization of resulting tissue models (Homan et al., 2019, Ko et al., 2019a, Shin et al., 2021, Sobrino et al., 2016). Although this remains moderately successful for the interpenetration of organoids, this strategy has been particularly promising for spheroid models.

The field of skin biology has seen the development of a range of skin equivalents interfaced with endothelial models. For example, endothelial barriers were formed at the bottoms or back of dermal compartments of human skin equivalents (Kwak et al., 2020, Lee et al., 2017, Wufuer et al., 2016) including in microfluidic chips, and micro-capillaries were allowed to assemble at the bottom of dermal constructs in transwell systems (Black et al., 1998, (Schurr et al., 1999) Such models of skin vascularization displayed improved epidermal barrier function (Kim et al., 2019, Kwak et al., 2020) and basal cell proliferation and differentiation (Kim et al., 2019), compared to non-endothelialised equivalents. However, few successful vascularized skin-equivalent models, integrated into microfluidic chips, have been reported to date.

In this report, we present the design of a full thickness human skin-on-a-chip model displaying a stratified differentiated epidermal compartment, assembled above a microvascularised dermal compartment (Figure 1). We first design a novel microfluidic chip featuring a circular central chamber separated from circumferential channels by micro-posts and characterize the impact of such structure on the morphology of the microvasculature. We then first identify culture conditions suitable for the formation of a stratified human skin equivalent in the resulting chips, prior to evaluating the impact of vascularization of the resulting tissue structure and epidermal differentiation. In addition, we introduce pericytes in the vascular compartment, in order to stabilize associated structures and evaluate the impact of such co-culture on the resulting vascularized skin equivalents. Overall, our results demonstrate the formation of vascularized skin-on-chip models that could be scaled and parallelized in order to design high throughput platforms for the testing of safety and efficacy of therapeutics.

**Figure 1.**
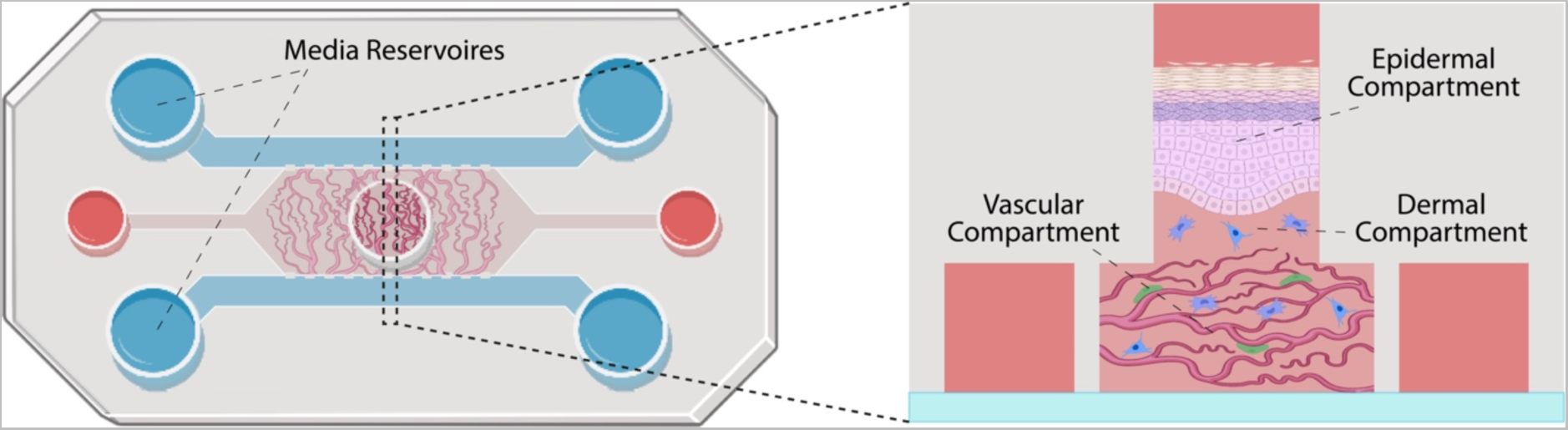
Schematic representation of the skin-on-a-chip model proposed. The chip is constituted of three parallel channels separated by microposts and connecting media reservoirs. A central well is positioned in the centre of the central channel. The vascular compartment is formed in the central channel (microvasculature shown in red and pericytes in green). Dermal fibroblasts (in blue) are introduced within a second hydrogel compartment, in the central well, before keratinocytes are seeded and allowed to stratify (in pink).

## 2 Materials and Methods

### 2.1 Chip Design and Manufacture

Chips were designed in AutoCAD by AutoDesk and printed on acetate sheets by Micro Lithography Services, Chelmsford, UK. Microfluidic chips were generated using Sylgard™ 184 PDMS and glass coverslips, following literature protocols (Jenkins, 2013). In short, masters were manufactured by coating clean silicon wafers with a negative photo resist (SU-2050) using a spin coater set to 500 rpm for 10 seconds (ramp 100 rpm/second for the initial 5 seconds) to distribute the photoresist, followed by a spin speed of 1700 rpm for 30 seconds (ramp 300 rpm/second for the initial 4 seconds) to spread the resist to a uniform depth of 80 μm. After an initial soft bake at 65°C for 3 min and then at 95°C for 10 min to harden the surface of the photoresist, photomasks corresponding to the desired chip micro-architectures were placed on the surface of the wafer and photocured with UV light at 90 mW/cm^2^ for 2.4 s, allowing resist crosslinking to occur. Resists were post-baked at 65°C for 3 min and then at 95°C for 10 min to assist the crosslinking, allowing to rest in between, then immersed in propylene glycol methyl ether acetate (PGMEA, Sigma Aldrich, 484431) for 5 minutes to dissolve un-crosslinked photoresist. Masters were immediately rinsed in isopropanol to remove excess PGMEA and dried under a stream of nitrogen. The resulting masters were examined using a reflected light microscope to confirm suitable patterning, and then incubated at 150°C for 3 min to hard bake the photoresist.. The PDMS base (Sylgard™ 184) was mixed with its curing agent at a 10:1 ratio, after which air bubbles were removed via centrifugation or under vacuum. The resulting formulation was poured onto the masters to a depth of 5 mm, placed under vacuum again to eliminated bubbles, and then left overnight at 70°C to allow full curing. The PDMS blocks were cut away from the masters to produce a stamp with features corresponding to the microarchitecture of the master. Biopsy punches were used to introduce cell culture medium reservoirs and hydrogel inlets, which can then be cleaned with residue free tape to remove any obvious debris and rinsed in deionized water and isopropanol or ethanol before drying in nitrogen. The stamps were placed in an air plasma system (HPT-200, Henniker Plasma) and treated for one minute with the channels facing upward, alongside glass coverslips. The PDMS blocks were then rapidly pushed firmly against the coverslip for 30 s to allow bonding and each chip was carefully inspected to ensure the system was fully sealed. Finally, chips were autoclaved and left in an oven for at least 48 h to allow the PDMS to recover its hydrophobicity.

### 2.2 Cell Culture

All primary cells used as part of this project were between passage numbers 3-6. Cells were cultured in T75 Nunc™ EasYFlask™ cell culture flasks (ThermoFisher®, 156499) at 37°C and 5% CO_2_ and passaged at approximately 80% confluency. HUVECs were obtained from Lonza™ and cultured in Endothelial Cell Growth Medium-2 (EGM-2, PromoCell® C-22111 or Lonza™ CC-3162). Human primary pericytes were purchased from PromoCell® (PromoCell®, C-12980) and cultured in Pericyte Growth Medium-2 (PGM2) also from PromoCell® (PromoCell®, C-28041). Detachment was achieved using a pericyte detachment kit (PromoCell®, C-41200), requiring addition of 5 mL HEPES buffered saline solution to the cells for washing, after medium aspiration, aspiration and addition of 5 mL Trypsin/EDTA 0.04%/0.03% for 3 min at room temperature. Trypsinisation was inhibited by addition of 5 mL trypsin neutralization solution, after which cells were centrifuged and resuspended in PGM2.

HCA2 fibroblasts - a human telomerase reverse transcriptase (hTERT) immortalized human dermal fibroblast cell line (Stephens et al., 2004) were cultured in Dulbecco’s modified Eagle medium (DMEM, Gibco™) supplemented with 10% fetal bovine serum (FBS, Gibco™ 26140079), 1% L-Glutamine and 1% penicillin-streptomycin (PS, Gibco™ 15140122). N/TERT cells (Hahn et al., 2000), a human keratinocyte cell line featuring constitutive expression of human telomerase reverse transcriptase (hTERT) were cultured in FAD medium (3:1 DMEM : Ham’s F12 Medium (ThermoFisher®, 11320074), supplemented with 1% PS, 1% L-Glutamine, 10% FBS, 0.5 µg/mL human hydrocortisone (Fisher Scientific®, 35245-0010), 8.47ng/ml cholera toxin (Sigma-Aldrich®, C8052-1mg), 10 ng/mL human epidermal growth factor (Peprotech®, AF-100-15) and 5 µg/mL human insulin (Sigma-Aldrich®, 15500).

### 2.3 Vasculogenesis Assays

A bovine fibrin gel was used as a matrix for vascularization, following protocols reported in the literature (Rohringer et al., 2014). In short, lyophilized bovine fibrinogen (Sigma-Aldrich®, F8630-25G) was dissolved in dPBS to a concentration of 20 mg/mL and filtered to ensure sterility. Separately, a bovine thrombin solution of 4 U/mL was prepared by dissolving lyophilized bovine thrombin derived from bovine plasma (Sigma-Aldrich, T4648-1KU) in 0.1% bovine serum albumin solution (BSA, Sigma-Aldrich, A9418-50G) in dPBS, which was added to EGM-2 to achieve the final concentration. Prior to injection into the device, 20 µL aliquots of fibrinogen solution were mixed with equal volumes of thrombin solution in which was suspended appropriate densities of HUVECs, which triggered formation of the final gel encapsulating endothelial cells.

For both chip designs, the wide central chamber was filled with 10 µL of a fibrin gel with a concentration of 10 mg/mL, in which HUVECs were suspended at a density of 6-10 million cells/mL by injection through the 3 mm central well with a cut 20 μL pipette tip. Both side channels were filled with EGM-2 supplemented with 50 ng/mL Vascular Endothelial Growth Factor (VEGF) through filling individual wells at one end of the chip and applying a negative pressure to the corresponding well for each channel using a 1000 μL pipette, therefore allowing channel perfusion. Medium reservoirs were filled equally. Chips were left to incubate at 37°C for approximately 45 min whilst HCA2 cells were passaged and encapsulated in a collagen gel at a density of 2 million cells/mL. Collagen gels were formed using rat-tail derived collagen I, as follows: working over ice, aqueous NaOH (1M) was added to 10X dPBS (ThermoFisher®, 70011044). Rat tail derived type I collagen (Corning®, 354249) was added to the buffer solution followed by HCA2 cells in dPBS and the mixture was well mixed, resulting in a final collagen concentration of 5 mg/mL and a HCA2 density of 2 million cells/mL. 30 μL of collagen gel solution containing fibroblasts was placed into the central well and allowed to set upon the HUVECs-loaded fibrin gel cast at the bottom. Medium wells and the central well were filled with EGM2 containing additional 50 ng/mL VEGF. Medium was replaced daily and after one week of culture were fixed in 4% PFA in PBS and stained for imaging.

### 2.4 Organotypic Skin-on-a-Chip Formation and Vascularization

#### 2.4.1 Dermal Equivalent preparation

Fibrin gels were prepared and injected into the vascular compartment of chips, either without cells, at 10^7^ cells/mL HUVECs or 10^7^ cells/mL HUVECs and 10^6^ cells/mL pericytes. 20 µL was injected into each chip. For vascularized chips, devices were filled with EGM-2 containing additional 50 ng/mL VEGF, which was replaced daily and cultured for 4 days before addition of the dermal and epidermal equivalents, to allow vascular development. Avascular chips were incubated with VEGF supplemented EGM2 24 h before addition of the skin equivalent. For embedding of the dermal equivalent gel precursor, solutions were stored on ice, using pre-cooled pipette tips kept in sterile sealed boxes at 0°C and microcentrifuge tubes to avoid premature Matrigel ® setting. Medium was removed from the reservoirs and central well of the devices. 250 µL high concentration (9 mg/mL) rat-tail derived collagen I solution was injected into a microcentrifuge tube, followed by 100 µL Matrigel ® (Corning®, 356234) and 50 µL 10X DMEM (ThermoFisher®, 11430030). After thorough mixing to ensure homogeneity, the pH was adjusted to 7 by dropwise addition of sterile 1 M NaOH, in aliquots of 5 µL, followed by further mixing, to a volume of approximately 405 µL. 50 µL sterile FBS was added and mixed in, and the mixture was put aside on ice until primary fibroblasts had been passaged and re-suspended to 1 M/mL in complete DMEM. 50 µL cell suspension was added to the collagen/Matrigel ® mixture and mixed thoroughly to a final fibroblast density of 10^5^ cells/mL. 20 µL of the resulting cell suspension in gel precursor mixture was injected into the central well of the device atop the fibrin scaffold. Injection was carried on the sides of the well, to avoid the formation of bubbles allowing the matrix to rest evenly across the surface of the fibrin gel underneath and to fill the entire diameter of the well.

#### 2.4.2 Epidermal Equivalent Preparation

Chips were incubated at 37°C for 1 h to allow gels to set, followed by passage of N/TERTs by addition of 10 mL sterile versene solution for 5 min followed by addition of 2 mL sterile 0.05% trypsin-EDTA for 5 min at 37°C. Cells were re-suspended to 10^6^ cells/mL in FAD and 25 µL of the suspension was injected on top of the gel down the side of the central well, to avoid bubble formation. After 10 min of incubation at room temperature to allow N/TERT attachment to the top of the dermal equivalent, the central well and medium wells were filled with FAD and chips were cultured overnight at 37°C and 5% CO_2_. After approximately 18 h of incubation, medium was aspirated from the chips and fresh FAD medium was replaced, which was done daily over the culture period. Chips were cultured for 7-14 days at 37°C and 5% CO_2_ before fixation in 4% PFA.

### 2.5 Imaging

#### 2.5.1 On-Chip Immunological Staining

For samples that were imaged on-chip, fixation was carried out in 4% PFA in PBS for 15 min, followed by washing in PBS thrice and incubated in 0.2% triton-X for 15 min. Primary and subsequently secondary stains were applied in 3% BSA overnight followed by a PBS wash. Chips were loaded whole into a sample holder mounted to the stage of the Zeiss™ LSM710 confocal microscope and imaged.

#### 2.5.2 Histology

##### 2.5.2.1 Cryo-Histological Sectioning

For sectioning, immunostaining and confocal imaging, after fixation, samples were immersed in 30% sucrose solution in PBS overnight by injection into lateral medium wells and the central well. This was aspirated the following day and replaced with a 10% sucrose/ 7.5% porcine gelatin solution in PBS and chips were left at 37°C for 15 min to allow the matrix to set. Simultaneously, some of the gelatin solution was carefully pipetted into well plates avoiding bubble formation to a depth of approximately 1 cm, and the plates were left at 4°C for 15 min to allow the matrix to set. Tissues were harvested from the central wells of the chips by slicing the coverslip from the PDMS carefully with a sharp scalpel and cutting chunks from the remaining block until the organotypic was exposed on 3 sides. At this point, the cultures were scooped from the PDMS and laid atop the set gels in well plates. More gelatin solution was injected to immerse the cultures completely and the well plates were placed back at 4°C for 15 min to set. A cold hexane bath was made by dropping pieces of dry ice into hexane in a deep glass dish with thick sides until it reached a temperature of between -30°C and -50°C. Orientation of the blocks was assured by cutting the corners of the blocks to be attached to the sample holder of the cryostat. Samples were frozen, sectioned and stored at -80°C until ready for staining.

##### 2.5.2.2 Paraffin Embedded Section De-Waxing and Antigen Unmasking

Redundant human skin was obtained from healthy donors following plastic surgery procedures at the Royal London Hospital or Phoenix Hospital, Essex. All donors provided written informed consent according to local ethical approval (East London Research Ethics Committee, study number 2011-000626-29 and East of England Research Ethics Committee, study number 21/EE/0057). Human skin sections embedded in paraffin wax were dewaxed by incubation at 55°C for 10 min to melt the paraffin followed by immersion in xylenes thrice, each for 15 min. To rehydrate the sections, slides were left in a bath of ethanol and xylenes at a 1:1 ratio for 5 min, followed by sequential incubation in 100%, 95% and 70% ethanol in deionized water, and finally pure deionized water (all twice, each for 2 min). To unmask antigens hidden during sample fixation, the deionized water was aspirated and sections were immersed in a 0.05% trypsin/0.1% CaCl_2_ solution in deionized water, pre-warmed to 37°C, and were incubated in a humid box for 20 min at 37°C. Samples were washed thrice more with dPBS before staining.

##### 2.5.2.3 Section Staining & Slide Preparation

Prior to staining, sections were allowed to equilibrate to room temperature for around 2 h. Sections were outlined with a hydrophobic pen to facilitate staining and reduce volumes of required reagents. 4% PFA in PBS was dropped onto each section for 15 min, followed by a PBS wash and further 15 min incubation with 0.2% triton-X in PBS. Following another PBS wash, sections were blocked overnight in 3% BSA solution in PBS at 4°C to reduce the risk of dehydration. Sections were incubated with 15 µL primary antibody solution overnight in 3% BSA at 4°C, according to the stain required. These included antibodies with the following antigens: keratin 14 (K14, AbCam® ab192055, 1:250), Involucrin (ThermoFisher®, MA511803, 1:40), Ki67 (RM9106-50, 1:125), laminin-332 (AbCam®, ab14509, 1:250), vimentin (AbCam®, ab8978, 1:150), TGM-1 (SigmaAldrich®, HPA040171, 1:150) and CD31 (BioLegend®, 303126, 1:100). Secondary antibodies were also applied overnight in a 3% BSA solution in PBS frequently alongside rhodamine-phalloidin and DAPI. These were Invitrogen® antibodies purchased from ThermoFisher® attached to AlexaFluor™ fluorophores with emission wavelengths of 488 nm or 647 nm. Secondary antibodies were raised against antigens according to the animal origins of the primary antibodies and diluted 1:1000.

Sections were washed with PBS which was carefully aspirated, followed by addition of Fluoromount-G™ Mounting Medium (ThermoFisher®, 00-4958-02) and a coverslip. Coverslips were allowed to attach at room temperature overnight and slides were carefully cleaned with ethanol prior to imaging. All fluorescent imaging was performed with the Zeiss™ LSM710 Confocal microscope.

### 2.6 Image & Statistical Analysis

For microvessel analysis, Z-stack files were loaded into Fiji and flattened to produce an image across the depth of the microvascularised compartment. These were analyzed in AngioTool (Zudaire et al., 2011) to find vessel junction number, overall vessel length and percentage area covered in any given image by vasculature. To analyze sections, Z-Stacks were taken through the entire depth of sections and loaded into Fiji. Individual slices were taken from the central image of each stack as representative of that stack and used for data acquisition and figure production. Statistical analyses were carried out using SPSS by IBM.

## 3. Results

### 3.1 Vasculogenesis-on-a-Chip

The proposed design of the microvascularised chip (Figure 1) is based on three channel chips previously reported by Kamm, Jeong and others (Hernández Vera et al., 2008, Jeon et al., 2014). In these microfluidic systems, three parallel linear channels are separated by micro-posts that can vary significantly in size and shape from one report to another. This enables the central channel to be loaded with an endothelial cell suspension in a gel precursor solution that is allowed to set *in situ* prior to filling the side channels with tissue culture medium. The device initially selected featured a hexagonal microfluidic chamber, 5 mm wide, with an integrated 3 mm diameter circular central well open to the air, allowing a minimal gap of 1 mm between the circumference of the well and the edge of the chamber. Peripheral medium channels were separated from the central chamber by 2 parallel series of hexagonal posts, 380 μm long and 90 μm wide, either side of the microfluidic culture compartment. The morphology and homogeneity of vascular networks formed in such systems were first investigated.

Vascularization was achieved via a vasculogenesis protocol adapted from the literature (Kim et al., 2016), consisting in de novo capillary formation of HUVECs encapsulated within the fibrin gel in the central compartment. The impact of cell density on vascular network formation was first examined (Supplementary Figure S1). HUVECs were introduced in the central gel compartment at 6, 8 and 10 million cells/mL. After 7 days of culture, cells formed interconnected vascular networks that were luminated, as evidenced by confocal microscopy (Figure 2). Cell densities had no significant impact on the morphology of the networks formed, within the range tested (Supplementary Figure S1). The total cross-section area of networks, the total length of capillaries and their degree of branching were comparable. However, we noted that the quality of the networks differed significantly depending on their position within the channel. Close to the central well (position A), vascular networks were denser (higher cross-section areas, length and branching densities) than in more distant areas, close to the injection channels (position B, Figure 2D-F).

**Figure 2.**
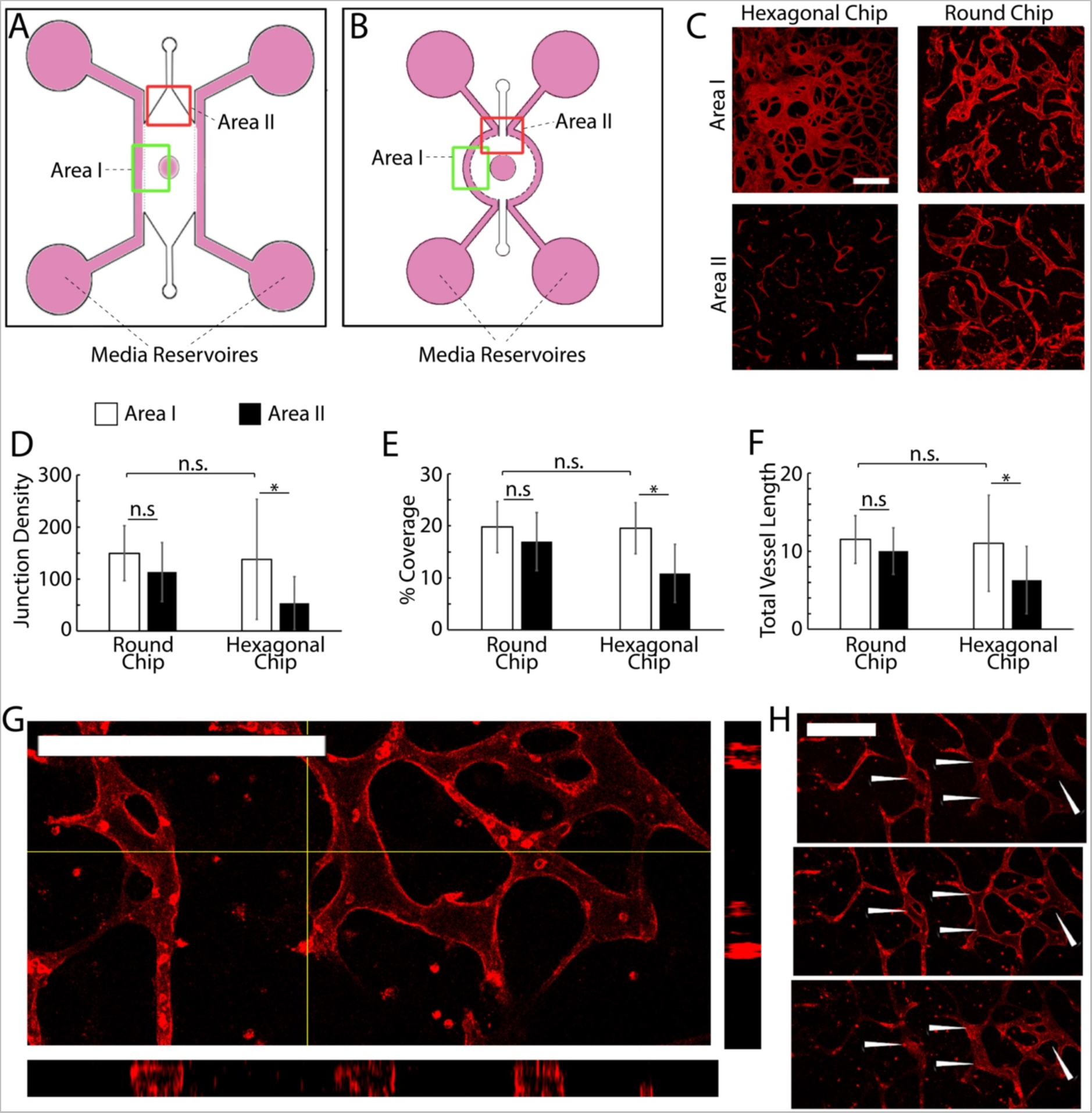
A, B) Schematic representation of the two chip designs investigated. C) Examples of microvascular networks formed in different areas of the central compartment of hexagonal and round chips. D-F) Quantification of the morphology of corresponding networks. G-H) Confocal images of networks formed in round chips displaying lumenated structures (G, z-stack projection and cross-section; H, corresponding individual sections taken across one network branch). *P < 0.05; n.s., non significant, P > 0.05; error bars are standard errors; n=3.

To encourage more uniform vascularization, the device was redesigned. The hexagonal microfluidic culture compartment flanked by straight parallel peripheral medium channels was replaced with a round compartment with circumferential 1 mm wide channel (Figure 2B). Hexagonal posts were redesigned to follow the circumference of the central channel. Given the lack of impact of HUVEC density on vascularization, within the range tested, cells were seeded within these redesigned chips at a density of 8 million cell/mL (suspended in 10 mg/mL fibrin gel).

Vascularization surrounding the central channel of round chips was comparable to hexagonal chips (Figure 2C-F), without significant changes in vascular network coverage, density or branching. In contrast to what had been observed with hexagonal chips, comparison of images taken near the central well of round chips with those taken near injection channels revealed no significant difference in network architecture either (Figure 2D-F). Levels of coverage by vascular networks were comparable near the central well and injection channels, and associated vascular networks were as dense and branched. This suggests that improved medium diffusion in different compartment within round chips enables improved formation of homogenous vascular networks. Examination of confocal z-stacks taken of corresponding vascular compartments also indicated the formation of a lumenated structures (20 µm of internal diameter) in networks formed in round chips (Figure 2G-H). Therefore our data indicate that round chips support the growth of a mature and more homogenous microvascular networks within multi-compartment microfluidic chips. This model was therefore adopted for the development vascularized skin-on-a-chip systems.

### 3.2 Establishment of a Skin-on-a-Chip Model

To engineer the full thickness dermal equivalent within our microfluidic model, we proposed to introduce dermal fibroblasts within the central well of round chips. As a first step, we introduced the dermal compartment on top of an acellular fibrin gel, which could subsequently be vascularized, as presented in Figures 1 and 2. A collagen I/Matrigel dermal equivalent matrix was selected, based on protocols established for transwell skin equivalent culture (Enjalbert et al., 2020). Fibrin was injected into the microfluidic compartment of the chip and devices were incubated at 37°C for 5 min, followed by perfusion of medium channels and filling of all on-chip wells with complete DMEM. Chips were left for 24 h at 37°C to allow bubbles forming at the medium-hydrogel interface to disperse. The medium was then removed from chips and 20 µL of dermal equivalent matrix (rat-tail derived collagen I and Matrigel with pH adjusted to 7) together with 10^5^ cells/mL primary human dermal fibroblasts were injected in the central well and allowed to set at the surface of the fibrin compartment, for 1 h at 37°C. N/TERT keratinocytes were passaged and resuspended at 1 million cells/mL in FAD medium, and 25 µL of the suspension was injected on top of the dermal equivalent. Devices were incubated for 10 min at room temperature to facilitate initial N/TERT attachment, followed by the filling of medium wells with FAD medium. After 24 h incubation, the medium was removed from the chip and replaced. To examine the impact of air-liquid interface culture, which is typically used for the formation of skin equivalents to encourage epidermal stratification and differentiation, dermal equivalents on chips were either left submerged (fresh FAD introduced into the central well) or exposed to air. Culture was continued for a total of 7 or 14 d, followed by fixation and embedding for histology and immunological staining.

At day 7, keratinocytes formed confluent epidermal layers across the surfaces of dermal matrices and positive for basal K14 (Supplementary Figure S2). After 14 days of culture, both in submerged conditions and at the air-liquid interface, we observed the formation of confluent stratified epidermal layers, atop the dermal equivalent matrix (Supplementary Figure S2). Infrequent proliferative cells (Ki67 positive) were observed in the basal epidermal layers of epidermal equivalents cultured submerged in medium (Figure 3). Although this localization and such overall low rates are typical of skin equivalents and proliferation rates observed in vivo (Heenen et al., 1998, Commandeur et al., 2012), the lack of Ki67 positive cells in tissues formed at the air-liquid interfaces was surprising. In agreement with such observations, we also observed significantly thicker epidermal equivalents in submerged cultures (46 +/- 16 µm) compared to those formed at the air-liquid interface (24 +/- 12 µm, Figure 3C). In addition, the abundance of the basement membrane protein laminin-332 was significantly higher in submerged cultures (by a factor of >2 fold, Figure 3C), suggesting that keratinocytes in these cultures were also more active at remodeling corresponding interfaces and recreating a normal functional basal lamina. The thickness of the basal marker keratin 14 was also significantly increased in submerged cultures (28 +/- 10 µm) compared to those kept at air-liquid interfaces (15 +/- 10 µm, Figure 4). Finally, whilst expression of the terminal differentiation marker involucrin was observed in the upper-epidermal layers of both cultures, the thickness of the associated differentiated layer was significantly increased in submerged cultures (3.5 +/- 2.2 µm) compared to those kept at the air-liquid interface (0.5 +/- 0.2 µm, Figure 4). However, relatively weak expression of the dermal fibroblast marker vimentin was observed in the dermal compartment of both cultures, and very weak staining for the epidermal differentiation marker transglutaminase 1 (TGM-1) was observed in the stratified layers of corresponding epidermal compartments (Figure 4B).

**Figure 3.**
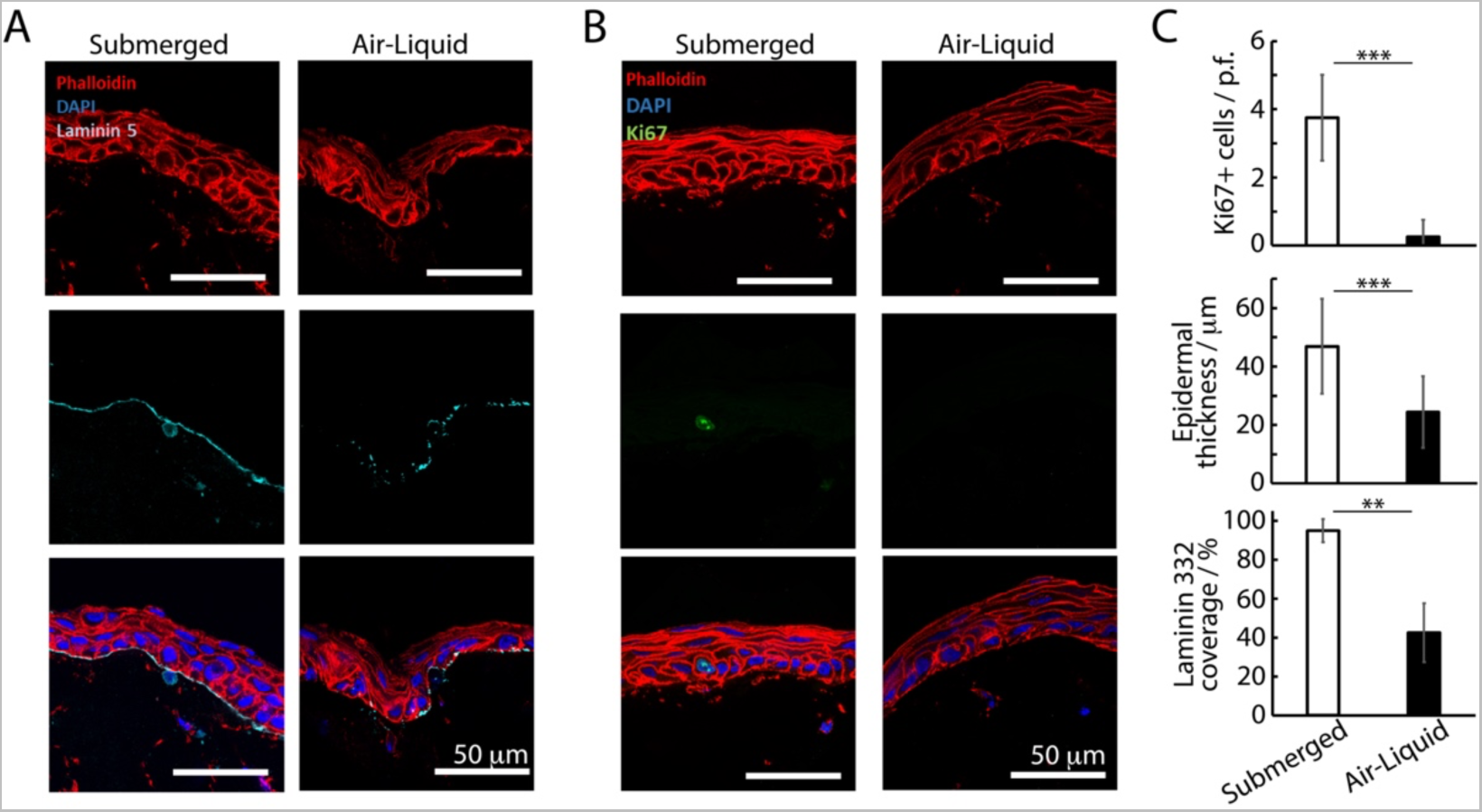
Histology sections of avascular skin-on-chip constructs immuno-stained with phalloidin (red), DAPI (blue) and for laminin 332 (A, cyan) or Ki67 (B, green). After an initial culture of 7 days, tissues were further cultured for 7 days in submerged or air-liquid conditions. C) Quantification of the corresponding density of Ki67+ cells, epidermal thickness and coverage of laminin 332. **P < 0.01; ***P < 0.001; n.s., non significant, P > 0.05; error bars are standard errors; n=3.

**Figure 4.**
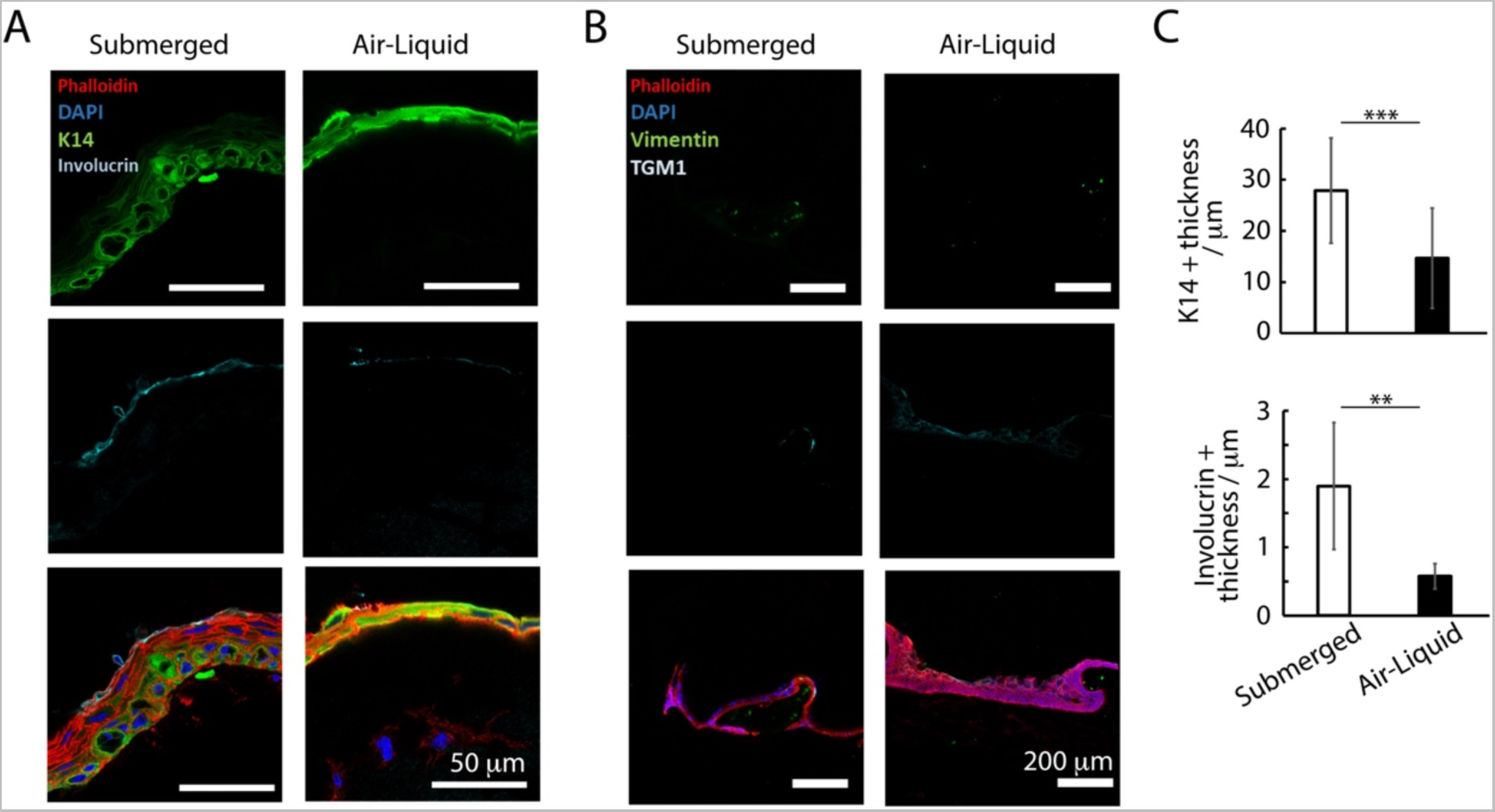
Histology sections of avascular skin-on-chip constructs immuno-stained with phalloidin (red), DAPI (blue), and for K14 and involucrin (green and cyan, respectively; A), or for vimentin and transglutaminase (green and cyan, respectively, B). After an initial culture of 7 days, tissues were further cultured for 7 days in submerged or air-liquid conditions. C) Quantification of the corresponding density of K14+ tissue thickness and involucrin positive tissue thickness. **P < 0.01; ***P < 0.001; n.s., non significant, P > 0.05; error bars are standard errors; n=3.

### 3.3 Microvascularised Skin-on-a-Chip Model

The microvascularised model developed in a fibrin matrix, and the stratified human skin equivalent were then combined in round chips. In addition, owing to their impact on the maturation and stabilization of microvasculatures (Campisi et al., 2018, Hirschi and D’Amore, 1996, Kim et al., 2013), we investigated the impact of pericytes on the resulting vascularized skin equivalents. HUVECs (10^7^ cells/mL) were seeded alone or together with human primary pericytes (10^6^ cells/mL) in a 10 mg/mL fibrin within the central channel of round chips. Side channels and the central well were supplemented with EGM-2 medium containing 50 ng/mL. After 4 days of culture to allow establishment of a mature microvasculature, the dermal equivalent was introduced into the central well, followed by seeding of N/TERT keratinocytes after 1 h, as previously described. Medium channels and the central well were filled with FAD medium and culture was continued for 14 d, under submerged conditions (Supplementary Figure S3). Cultures were fixed and dissected from chips for histological processing and immunological staining. Sections taken from our samples were compared with *ex vivo* human skin sections.

Resulting tissues displayed a well-structured multi-layer epidermal equivalent with flattened stratified squamous cells (Figure 5 and Supplementary Figures S4 and S5). Although the overall thickness and level of flattening and stratification of the epidermal compartment remained lower than what can be observed in sections from human skin, epidermal equivalents were more developed in the presence of microvasculatures (whether with or without pericytes). Rete ridges present in *in vivo* human skin were also not observed in our HSE-chip models, although these are not generally recapitulated in other human skin equivalent reported (Choudhury and Das, 2021). The overall depth of the epidermal compartment significantly increased in vascularized chips (86 +/- 20.3 µm, for vascularized chips in the presence of pericytes, compared to 64 +/- 26.5 µm, for avascular chips; Figure 5C). Although these thicknesses are in line with those reported for skin equivalents (Chopra et al., 2015, Wei et al., 2017), our results indicate that vascularization impacts on the establishment of the epidermal structure in skin equivalents.

**Figure 5.**
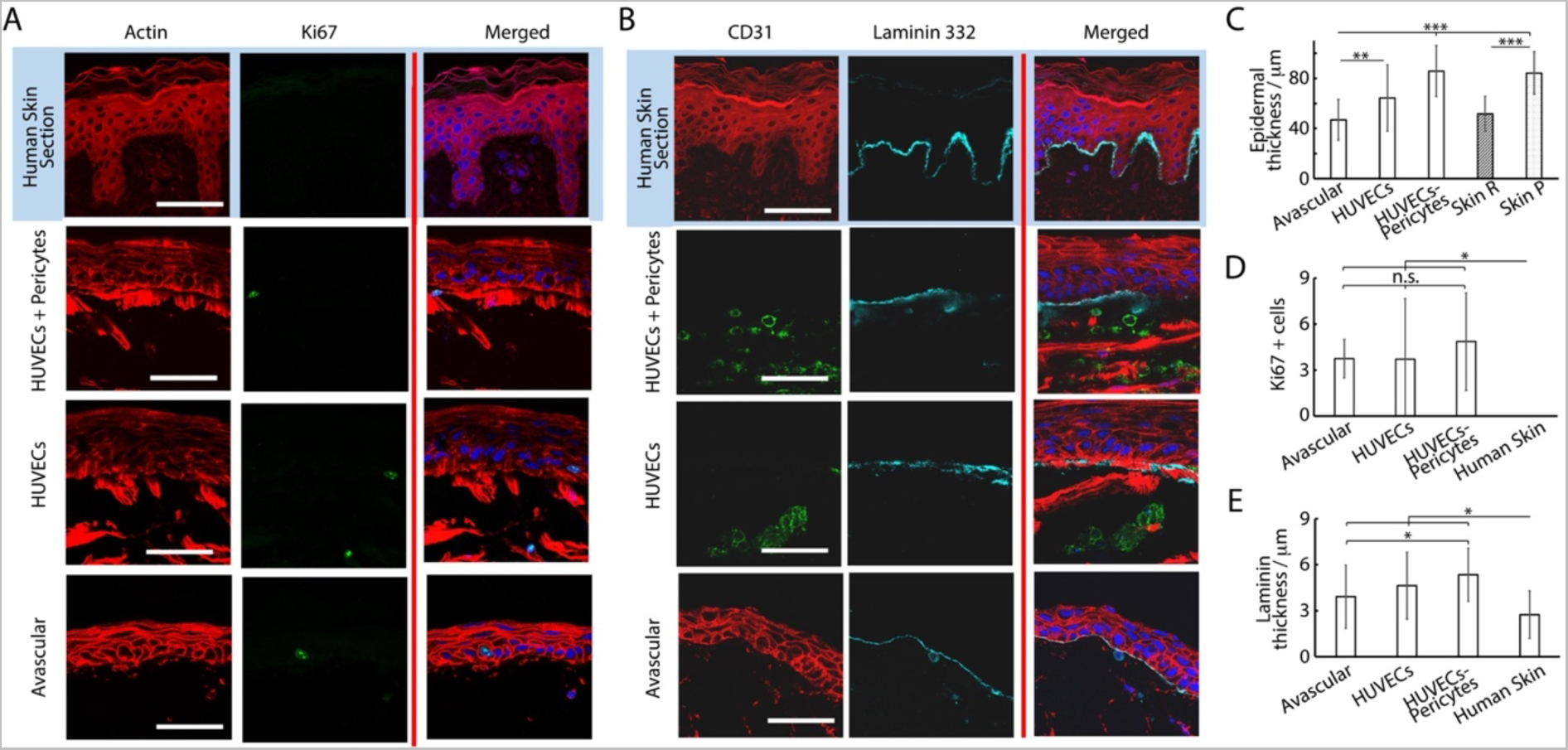
Histology sections of microvascularised skin-on-chip constructs and human skin sections immuno-stained with phalloidin (red), DAPI (blue) and for Ki67 (A, green) or CD31 and laminin 332 (B, green and cyan, respectively). Tissues were cultured for 14 days in submerged conditions. C) Quantification of the corresponding epidermal thickness (C), density of Ki67+ cells (D) and thickness of laminin 332 (E). *P < 0.05; **P < 0.01; ***P < 0.001; n.s., non significant, P > 0.05; error bars are standard errors; n=3.

To study this phenomenon further, we investigated stainings of tissue sections. The impact of vascularization on the basal compartment of the epidermal equivalent was relatively modest. We observed a small increase in the thickness of the basement membrane constituent laminin 332 in vascularized chips, compared to avascular models and *ex vivo* skin sections (Figures 5B and E). This increase was not statistically significant when comparing avascular and vascularized chips, but P values decreased to <0.001 when pericyte/HUVEC vascular networks were used in the vascularised compartment, compared to human skin sections. This modest change in basement membrane did not translate into differences in expression of the basal marker keratin 14 (Figure 6). Similarly, Ki67 positive cells in the epidermis were confined to the basal layer in all conditions and their frequency was comparable (Figure 5D). However, these observations contrasted with the lack of Ki67 expression in the basal compartment of *ex vivo* human skin sections, in line with previous observations made on comparable human sections (Heenen et al., 1998).

**Figure 6.**
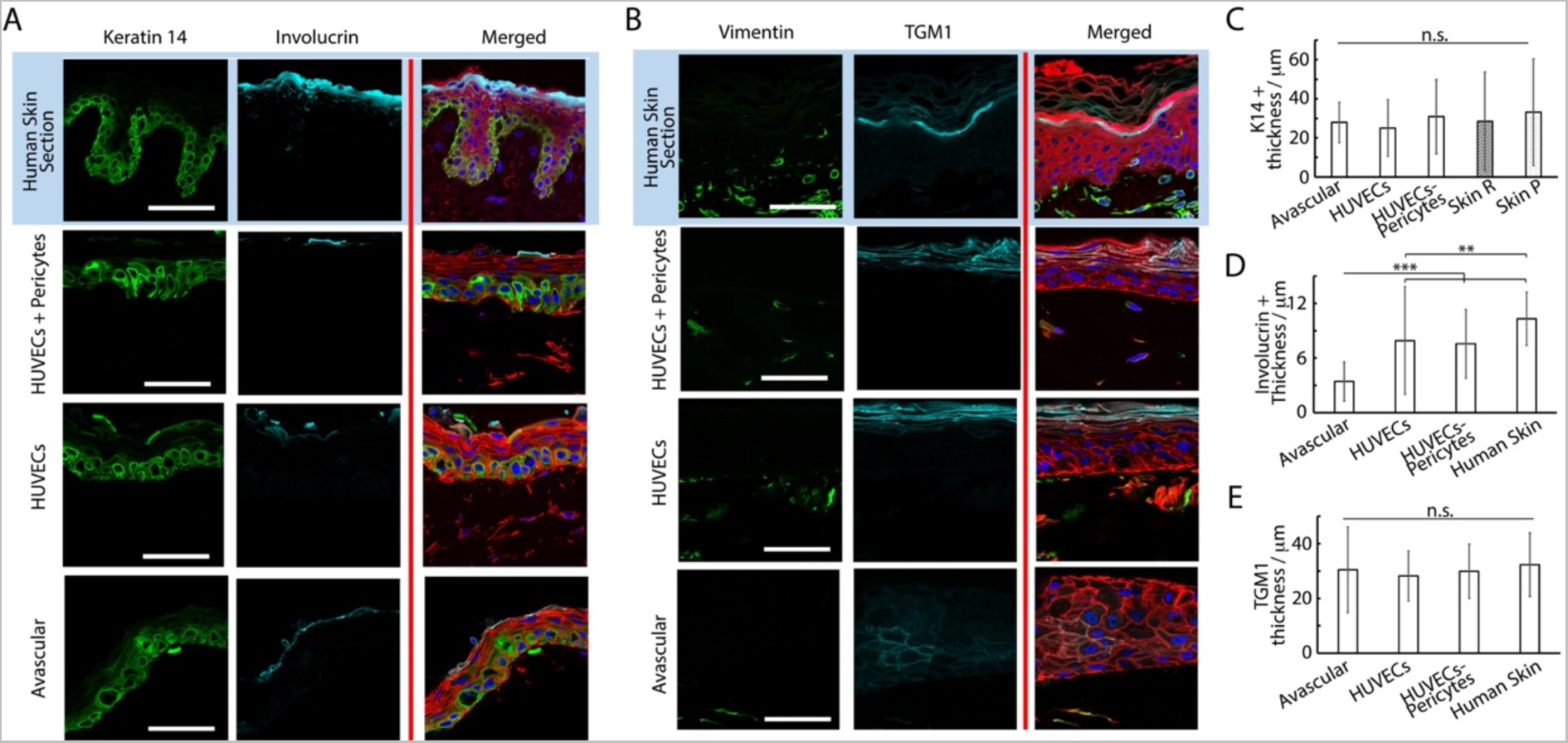
Histology sections of microvascularised skin-on-chip constructs and human skin sections immuno-stained with phalloidin (red), DAPI (blue) and for K14 and involucrin (A, green and cyan, respectively) or vimentin and transglutaminase (B, green and cyan, respectively). Tissues were cultured for 14 days in submerged conditions. C) Quantification of the corresponding K14+ tissue thickness (C), involucrin + tissue thickness (D) and transglutaminase + tissue thickness (E). **P < 0.01; ***P < 0.001; n.s., non significant, P > 0.05; error bars are standard errors; n=3.

Epidermal differentiation marker expression was more established and consistent with the addition of a vascularized dermal compartment: the thickness of involucrin positive layers in chips containing either HUVECs alone or alongside pericytes (8 +/- 5.8 and 8 +/- 3.8 µm, respectively) compared to avascular models (3 +/- 2.2 µm, Figures A and D). However, involucrin expression remained weaker than in human skin sections (10 +/- 2.9 µm). Similarly, staining for transglutaminase was significantly more pronounced in vascularized, compared to avascular chips. Beyond expression levels, the localization of transglutaminase was also more obviously localized at the upper layers of the epidermis in vascularized HSE-chips, recapitulating well the pattern of expression typical of human skin sections (Figure 6B). We did not observe a significant impact of vascularization on the thickness of transglutaminase expression (Figure 6E), but expression was not confined to the uppermost layers of the epidermal equivalents in avascular chips and could be seen in layers directly above the basal layer.

The impact of vascularization on the dermal compartment could also be clearly observed. CD31 positive structures were obviously only observed in vascularized models, as would be expected from the presence of endothelial cells (Figure 5B). Associated with such structures, we clearly observed an increase in vimentin positive cells in vascularized chips, compared to avascular cultures (Figure 6B). Vimentin positive cells were clearly denser in human skin histological sections, but this reflects a general higher degree of cellularity in normal tissue dermis, compared to the in vitro models established.

Further characterization of the vascular compartment was carried out next. We observed extensive dermal microvascular networks spanning the microfluidic compartments underlying the central well cultures of chips loaded with HUVECs alone (bright field images, Figure 7), confirmed via staining for the endothelial marker CD31 (Figure 7D). In the presence of pericytes, vascular networks were also clearly observed, but displayed altered morphologies, in agreement with similar systems reported in the literature (Kim et al., 2015b). Pericyte co-cultures displayed reduced total length and branching (Figures 7A-B). Such phenotype is proposed to be associated with the maturing and stabilizing role of pericytes, enabling to better retain the stability of resulting networks. Mechanisms involved in this behavior remain partially unclear, but may involve cytokine secretion (e.g. angiopoietin-1, angiogenin, HGF, TGF-a and TNF) and perivascular matrix remodeling (Newman et al., 2011). Whilst others reported an impact on endothelial cell-cell junctions (Jeon et al., 2014, Jeon et al., 2015), our previous results did not confirm such effect.

**Figure 7.**
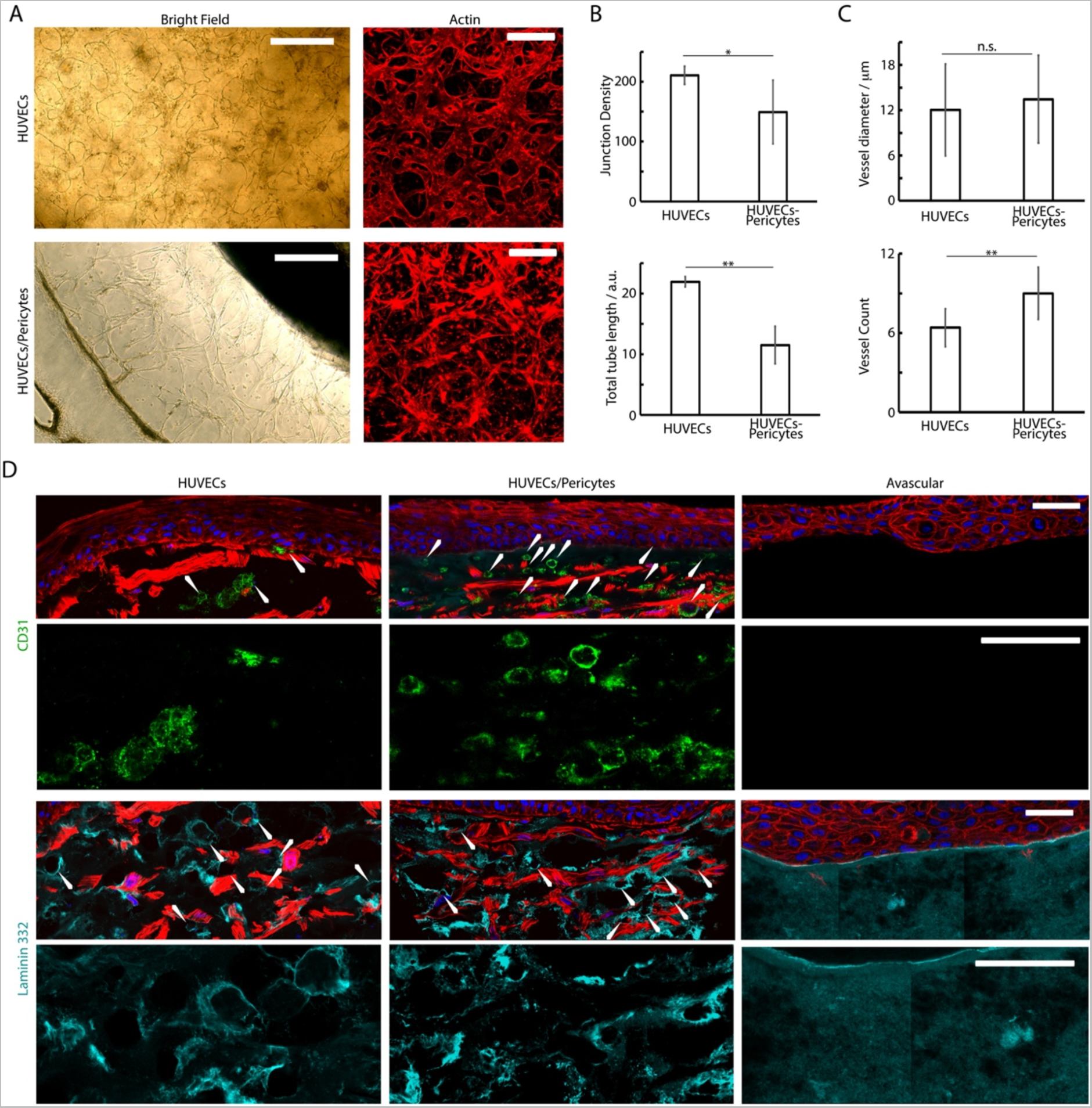
A) Brightfield and confocal images (phalloidin) of the basal microvascular compartment of chips injected with either HUVECs alone or HUVEC and pericytes, followed by 4 days of culture in EGM2 medium + 50 ng/mL VEGF and 2 weeks in FAD medium. B and C) Characterisation of the morphology of the corresponding networks. D) Histology sections of microvascularised skin-on-chip constructs immuno-stained with phalloidin (red), DAPI (blue), and for CD31 (green) and laminin 332 (cyan). Tissues were cultured for 14 days in submerged conditions. *P < 0.05; **P < 0.01; n.s., non significant, P > 0.05; error bars are standard errors; n=3.

To investigate the architecture of the microvasculatures integrated into the dermal equivalents in our cultures, we examined the formation of capillaries in the sections generated (Figure 7D). CD31 and laminin stainings clearly established the formation of rosettes associated with the formation of lumenated capillaries (Figure 7D), in agreement with the recruitment of laminin at the basal membrane of micro- capillaries (Black et al., 1998, Schurr et al., 1999, Kubota et al., 1988). The density of these structures was clearly increased in HUVEC-pericytes co-cultures, indicating a better retention of the vascular networks within these models (Figures 7C and D). The average external diameter of capillaries determined from sections, although approximative as only relying on the ability to identify clear transverse sections, were 12 +/- 6 µm for monocultures and 13 +/-6 µm for HUVEC/pericytes co-cultures, which correspond to the external diameter of dermal arterial capillaries *in vivo*, identified from human skin sections (Braverman, 2000). Overall, our results are evidencing the supportive role played by pericytes in our model, maintaining the microvasculature integrity despite the use of mixed culture media in our protocol (Campisi et al., 2018, Hirschi and D’Amore, 1996).

## 4. Discussion

Altogether these data demonstrate that our chip design enables the formation of a microvascular network embedded in a multi-compartment microfluidic system and underlying the formation of a dermal/epidermal skin. The design of a round central chamber and circumferential channels enables the formation of a more homogenous microvasculature within the central compartment (Figure 2), in agreement with previous data presented for a vascularized ocular model (Ko et al., 2019b). It is also expected that the circular design selected will result in more homogenous cytokine and growth factor gradients within the different compartments, as was previously described in a model of pluripotent stem cell differentiation (Manfrin et al., 2019) and as briefly discussed in a model of central nervous system axonal growth, across the diameter of a microfluidic compartment (Park et al., 2009).

A surprising result from our study is that skin equivalents, in the absence of microvasculature, form more mature stratified epidermal compartments in submerged conditions, compared to those cultured at the air-liquid interface. This contrasts with the literature indicating that culture at the air interface, for skin equivalents grown in transwell systems, allows improved stratification and accurate epidermal differentiation marker expression (Prunieras et al., 1983, Rosdy and Clauss, 1990). These results may be explained by the smaller dimensions of the constructs generated and the lower volumes of media required for their culture. Indeed, the smaller dimension may result in higher local oxygen concentrations, impacting on keratinocyte commitment and stratification (Ngo et al., 2007). In addition, the smaller volumes may allow greater accumulation of cytokines and growth factors promoting stratification. It may also be that, the lower volumes of media used in culture at air-liquid interfaces in our chips result in increased cell death, and that the added volume associated with submerged culture helps limiting such process, as demonstrated by Lee et al. in 2017 in their vascularized skin-on-a-chip (Lee et al., 2017). Indeed, tissues formed at the air-liquid interface displayed significantly reduced Ki67 expression, compared to submerged systems (Figure 3).

Culture on microvascularized bed further improved the stratification and differentiation of skin equivalents. The use of pericytes was found to have weak but statistically significant impacts on the density of capillaries and their apparent lumenation in our skin equivalents (Figure 7). Therefore, beyond their initial impact on the morphology of vascular networks, pericytes may contribute to the complementarity of vascularized models with other tissue equivalents, allowing the formation of more complex vascularized tissue models. This may be associated with perivascular matrix remodeling (Avolio et al., 2017, Bergers and Song, 2005) and cytokine secretion, which were found to promote the stability of vascular networks (Hurtado-Alvarado et al., 2014, Sounni et al., 2011).

Improved epidermal stratification and differentiation upon addition of a microvascular component has previously been demonstrated in both *in vivo* mouse and *in vitro* systems (Baltazar et al., 2020). Conversely, Rochon et al. have shown that various epithelial cell types have a direct impact on endothelial vessel physiology *in vitro* (Rochon et al., 2009). This is likely due to increased bi-directional diffusion of secreted growth factors and cytokines in the surrounding microenvironment, known to impact differentiation in keratinocyte populations (Ansel et al., 1990, Gröne, 2002). Stromal fibroblasts in the dermal equivalent matrix also likely contribute to the secreted factors to encourage improved epidermal stratification; this has been demonstrated previously in human *in vitro* systems, which commonly report an undifferentiated, unstratified epidermis’ with a reduced thickness and poor localization of upper-epidermal and basement membrane markers in the absence of proximal fibroblasts (Saintigny et al., 1993). In 2020, Russo et al. published an extensive review on this topic (Russo et al., 2020). Pericytes alongside HUVECs have been shown to increase epidermal thickness and the expression of cytokeratins 1/10 and laminin 332 in *in vitro* skin models, which was also confirmed in engraftment experiments into a murine setting for a duration of 4 weeks (Baltazar et al., 2020). In agreement with this literature, stromal cell interactions are proposed to improve epidermal architecture in our model.

Overall, our results demonstrate the feasibility of hierarchically structured vascularized-skin-on-chip equivalents, paving the way for the embedding of these systems for testing of therapeutics safety and efficacy testing in a human context in vitro. Such models will allow testing of systemic as well as topical delivery in realistic scenarios, in platforms that may be compatible with high throughput testing. This still requires the engineering of multi-chip plate systems compatible with high throughput formats, in analogy to some of the epithelial/endothelial tissue models developed on the Emulate platforms (Si et al., 2021), and the parallelization of multi-cellular culture and embedding in the resulting micro-engineered chips. This latter process may require particular attention from the research community as parallelized and automated multi-cell culture systems have not been engineered to date.

## 4 Author Contributions

CFEJ, JEG and JC: conceptualization and project administration. CFEJ: data curation, formal analysis, and validation, visualization and writing – original draft. JEG and JC: funding acquisition, resources and supervision. SC: data curation. JEG: writing – review and editing. All authors contributed to the article and approved the submitted version.

## 6 Acknowledgments

The authors would like to acknowledge the contribution of Atiya Sarmin (Blizard Institute, Whitechapel) for helping to formulate the dermal equivalent employed in our model.

## 7 Supplementary Material

Supplementary immunostainings of vascularised chips and histology sections are available as supplementary information or from the authors.

## 8 Data Availability Statement

The raw data supporting the conclusions of this article will be made available by the authors, without undue reservation.

