## Supplementary information for "Design of an Integrated Microvascularised Human Skin-on-a-Chip Tissue Equivalent Model"

**Christian F. E. Jones[1,2], Stefania di Ciò[1,2], John Connelly[3], Julien Gautrot[1,2*]**

[1]Institute of Bioengineering and [2]School of Engineering and Materials Science, Queen Mary University of London, London, United Kingdom

[3]The Blizard Institute, Queen Mary University of London, London, United Kingdom

*** Correspondence:**
Julien Gautrot


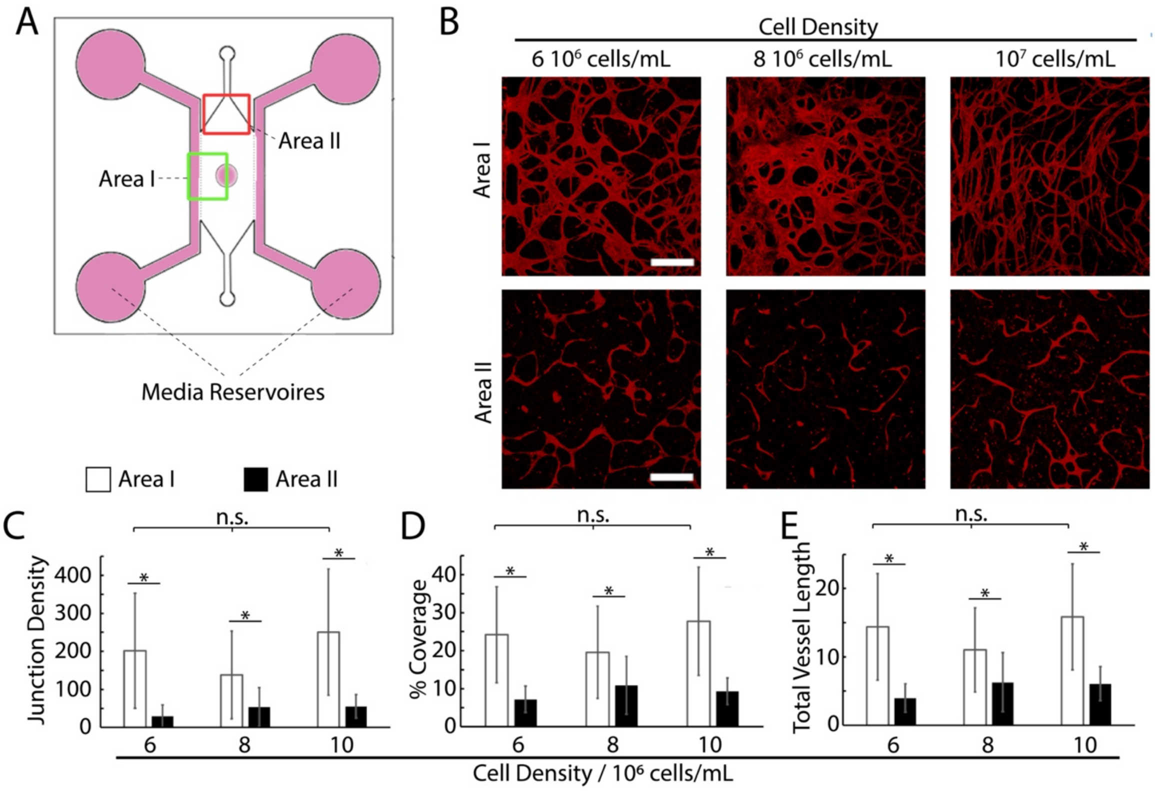


**Supplementary Figure S1.** Impact of cell density on vasculogenesis. HUVECs were seeded in fibrin gels (10 mg/mL) at densities of 6, 8 and 10 million cells / mL. A) Schematic representation of the chip used for this experiment. B) Confocal microscopy images of resulting networks (CD31, red). C-E) Quantification of the morphology of corresponding networks. *P < 0.05; n.s., non significant, P > 0.05; error bars are standard errors; n=3.

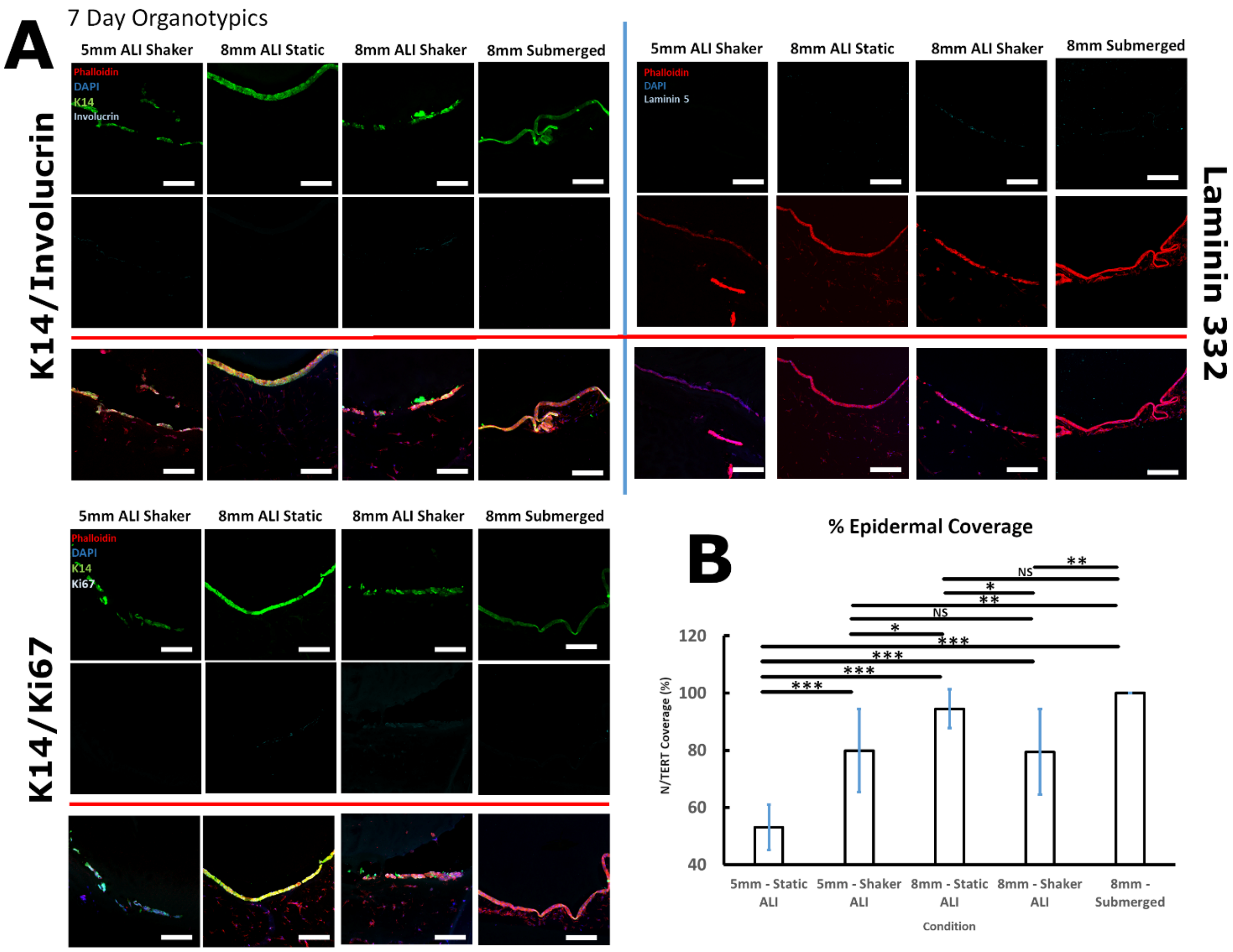


**Supplementary Figure S2.** A) Histology sections of avascular skin-on-chip constructs immuno-stained with phalloidin (red), DAPI (blue) and for K14, involucrin, laminin 332 and Ki67 (see figure for details of color channels associated with each marker). Cultures were incubated for 24 h following N/TERT seeding to allow adhesion, followed by 6 days at air-liquid interfaces (ALI) or submerged. Two well diameters (5 and 8 mm) were investigated. Chips were either incubated in static conditions or on an obrbital rocker. Medium was changed daily. B) Quantification of the corresponding density of keratinocyte coverage. *P < 0.05; **P < 0.01; ***P < 0.001; n.s., non significant, P > 0.05; error bars are standard errors; n=3.

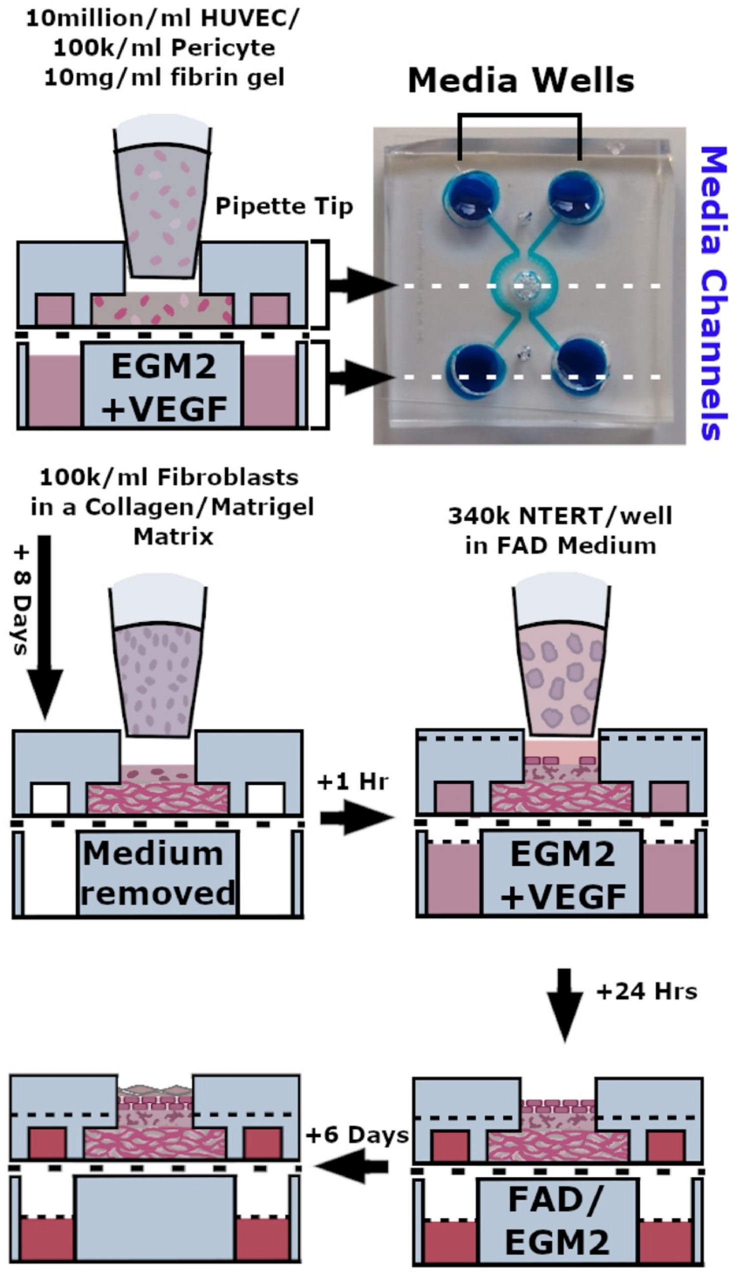


**Supplementary Figure S3.** Schematic representation of structure of the skin-on-a-chip model proposed and the different steps associated with their formation.

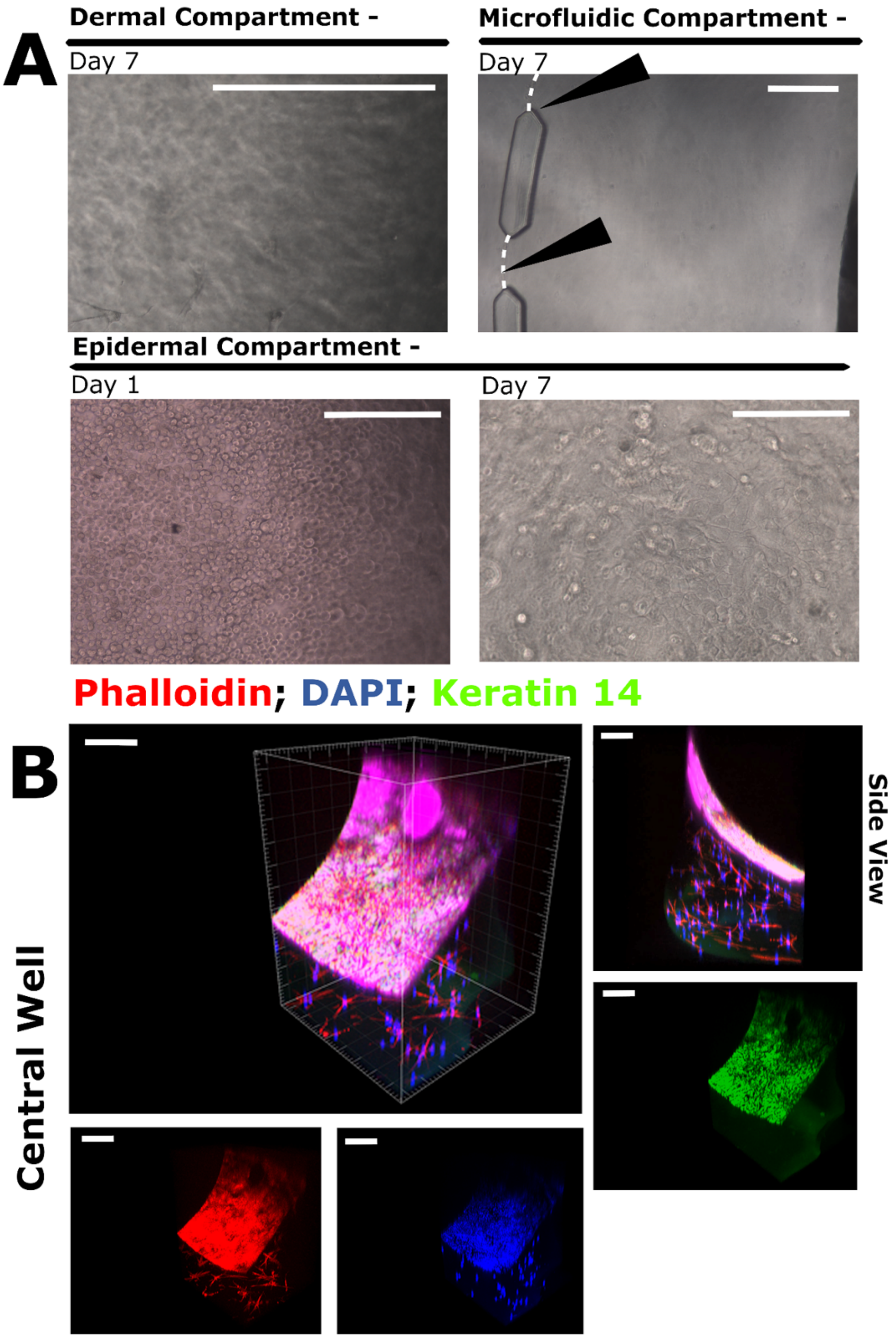


**Supplementary Figure S4.** BF images (A) and confocal microscopy images (B) of our skin-on-a-chip models. A) After 7 days of culture at the air-liquid interface, the dermal component featured fibroblasts with a spread morphologies and the epidermis displayed a confluent cobblestone architecture. The fibrin peripheral compartment had not contracted away from the PDMS posts and hence a hydrogel-medium interface was maintained (white dotted line, black arrows). B) Immunostaining and confocal images presenting the structure of the dermal-epidermal constructs and confirming the formation of a confluent K14 positive epidermal layer and an underlying dermal equivalent compartment, populated with fibroblasts. Scale bars: 200 µm

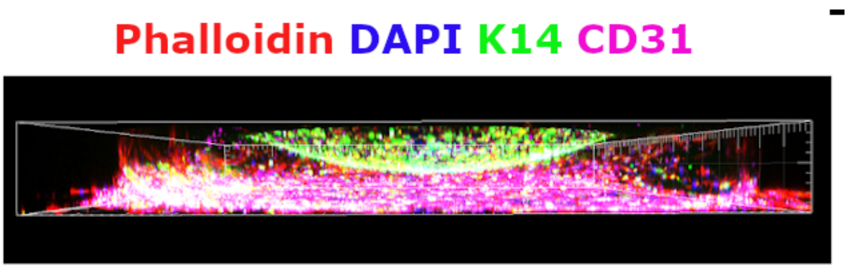


**Supplementary Figure S5.** Confocal microscopy images presenting the structure of the dermal-epidermal constructs.

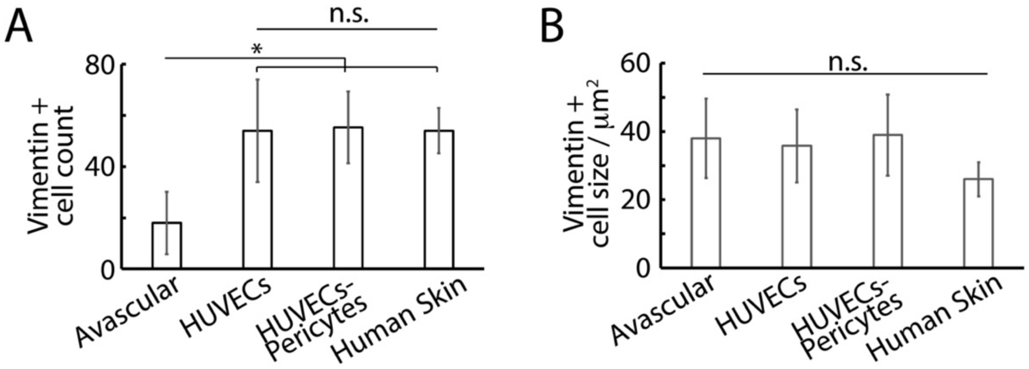


**Supplementary Figure S6.** Quantification of the vimentin + cells, from hisology sections of vascularised skin-on-chip constructs and human skin sections (tissues were cultured for 14 days in submerged conditions). *P < 0.05; **P < 0.01; n.s., non significant, P > 0.05; error bars are standard errors; n=3.
